# Precision Medicine in the Clinic: Personalizing a Model of Glioblastoma Through Parameterization

**DOI:** 10.1101/037424

**Authors:** Andrea Hawkins-Daarud, Kristin Swanson

## Abstract

In this article, we present an example of how to parameterize a partial differential equation model of glioblastoma growth for individual patients. The parameters allow for a deeper understanding of individual patients which have all been labeled with the same disease based on the tumor growth kinetics.

## I. Glioblastoma in the Clinic

Glioblastoma is a deadly type of primary brain tumor. It is known that the cancer cells are diffusely present throughout the brain, but the degree of invasiveness is heterogeneous throughout the patient population. Unfortunately, there is not a currently accepted method to identify for each patient the extent of the tumor cells. Clinically, this results in patients being treated in a very aggressive, one-size-fits-all manner.

Due to their location in the brain, these tumors are primarily monitored by magnetic resonance images (MRI). The main sequences that are used are the T1-weighted with gadolinium contrast enhancement (T1Gd) and T2-weighted sequences. It is commonly accepted that abnormalities on the T1Gd image highlights tissue primarily composed of tumor cells and the abnormalities on the T2 image highlight regions of lower tumor cell density and swelling. However, there is no way to determine the full extent of the tumor cells.

In general, treatment for patients with glioblastoma begins with a surgeon trying to remove as much of the abnormality seen on the T1Gd as possible. As this surgery is quite invasive, the merits of such an aggressive surgery have been debated [1], [2]. However, the current consensus in the field, based on recent studies, is that this aggressive removal of tissue results in longer survival [3]. These studies focused on the population level rather than the individual.

## II. Model Parameters Identify Glioblastoma Patients Who Will Benefit from Surgery

Over the past decade, our group has harnessed a partial differential equation model, representative of glioblastoma growth, to try and glean insights on the heterogeneity of the disease and what it means for individual patients [4]–[7]. This model captures only the basic definition of cancer in that the cells are known to proliferate and to invade. Given just a few assumptions of how the MRI abnormalities relate to the tumor cell density (described more in the methods section), we have been able to estimate net rates of proliferation and invasion for individual patients. These parameters can be used to estimate the nodularity of an individual tumor, that is, how much of the tumor is actually visible on the MRI. We hypothesized that the patients that were defined as more nodular by our model would receive greater benefit in terms of survival from an aggressive surgery trying to remove all of the abnormality on the MRI.

**Figure 1.**
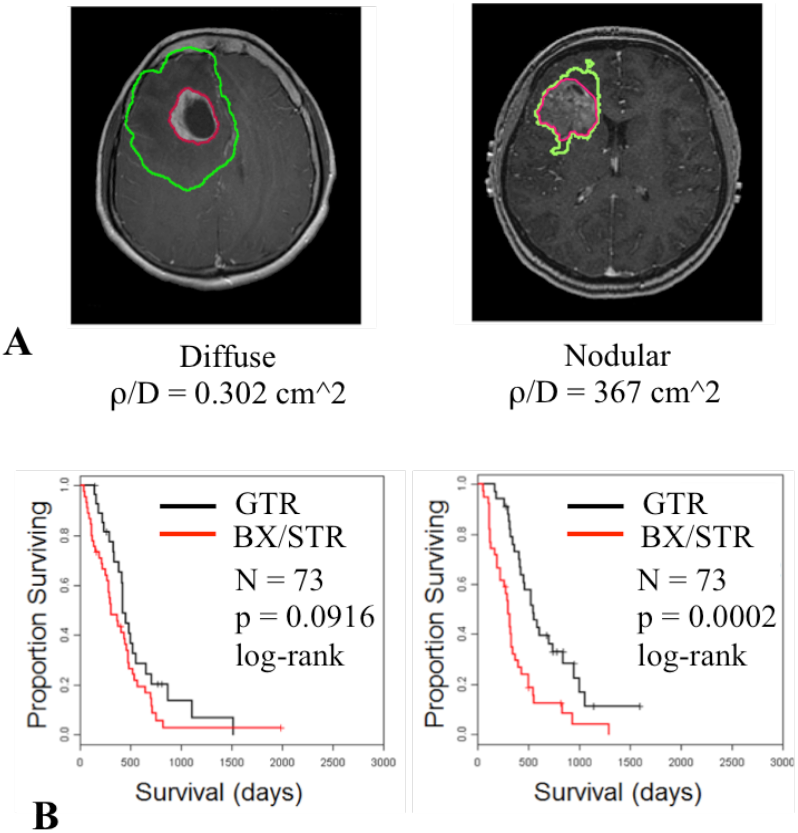
A) Examples of patients classified as having a diffuse (left) and nodular (right) tumor burden. The red line highlights what portion of the brain the surgeon would try and remove for a gross total resection and the green line highlights the model predicted region containing 99% of the tumor cells. B) Survival curves for the two identified patient populations (diffuse on left, nodular on right) demonstrating the difference in survival between those receiving a gross total resection (GTR) and a biopsy (BX) or subtotal resection (STR) only. Only in the nodular patients is a statistically significant difference observed.

To investigate our hypothesis, we retrospectively identified 243 contrast enhancing glioblastoma patients from our IRB approved database receiving a gross total resection (GTR), a subtotal resection (STR), or a biopsy (Bx) only. We then estimated our model defined net rate of proliferation and invasion based on imaging for each of the patients and separated them into three groups. Patients with nodular tumors, patients with diffuse tumors, and patients in between. Within each cohort, we then identified those that had an aggressive GTR versus those with only a STR or Bx. By comparing the survival curves within each nodularity group between the GTR patients and the STR/Bx patients, we were able to show that the only case where there was a significant difference in overall survival was for the nodular tumor group. These results were published in [8]

## III. Parameterizing Partial Differential Equation Models

### A. Partial Differential Equation Models

Partial differential equations (PDEs) are equations including multivariable functions and their partial derivatives. It is often the case that the partial derivatives are with respect to time and spatial dimensions. This is particularly true for partial differential equations representative of physical phenomena. Mathematical models based on PDEs generally must be augmented with initial conditions and boundary conditions. In a few cases, the functions solving such models can be solved for analytically but more often numerical methods are utilized to find approximate solutions [9]–[11].

The specific model used for the presented results is a relatively simple model capturing the basic definition of cancer: cells proliferate uncontrollably and they invade the surrounding tissue. Mathematically, this can be written as:

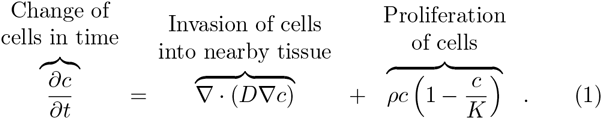

Here *c* is a multivariable function of time and space representative of tumor cell density evolving over time. To complete the model, this equation is augmented with Neumann boundary conditions representative of cells not being able to leave the domain and a point source initial condition or one more reflective of a particular patients disease burden. It can be solved as a three-dimensional spherically symmetric problem or in an anatomically accurate setting representative of the brain.

This model is general, but can be made specific to an individual patient by specifying the two parameters *D* [mm^2^/yr], the net rate of invasion (also known as diffusion), and ρ [1/yr], the net rate of proliferation. *K* is the carrying capacity and is assumed to be constant for all patients.

### B. Calibrating Parameter Values

In general, PDE models are applicable in different scenarios (i.e. prediction of two different tumors’ growth patterns) by modifying key parameters. The process of finding the best parameters to match a particular scenario is referred to as parameter calibration and requires data representative of the scenario of interest for model prediction comparison. Determining the best data with which to calibrate can be a challenge, however, often the data available is limited which means the greater challenge is how to use the data available to calibrate the model

For (1), under reasonable assumptions regarding the initial condition, the solution is known to set up a traveling wave, that is the function that solves the equation has the same shape that moves outward in time [12] (Chapter 11). The shape is uniquely determined by the parameters *D* and ρ. Further, it can be shown that in one-dimension this wave moves precisely at the radial velocity of 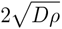 [12] (Chapter 11). In spherical symmetry, an analytic solution sadly does not exist. However, as predicted tumor size increases, the solution tends asymptotically to a travelling wavefront with a radial velocity 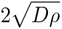.

**Figure 2.**
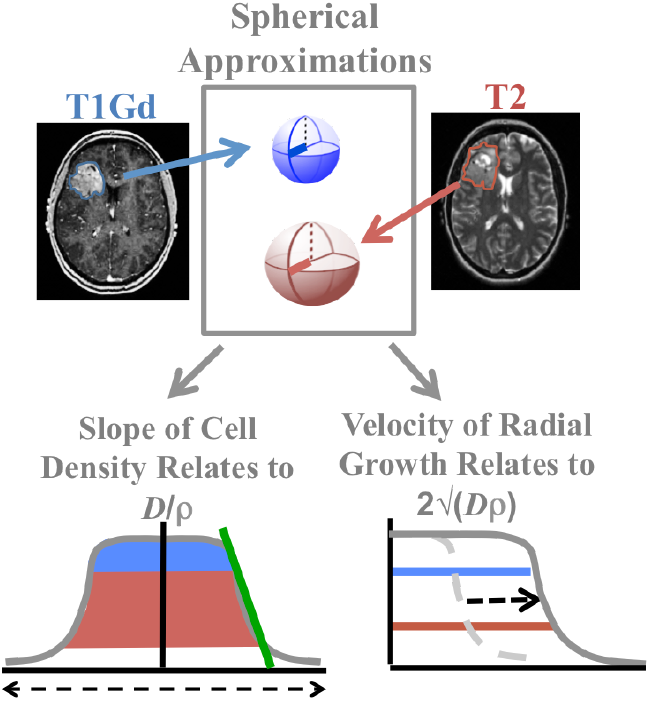
*Process of data utilization for parameter calibration of the model for glioblastoma growth.* Measured abnormality volumes on MRIs are converted to spherically equivalent radii. By using a T1Gd and T2 radius from the same day, the slope of the cell density can be estimated providing a value for D/r. Using either T1Gd or T2 images from two time points, a radial velocity can also be estimated providing a value for 2√(D_ρ_).

Despite the non-spherical shape of many glioblastomas seen on MRIs, we have found that considering patients’ tumors in this way is still very instructive. Thus, for individualizing the two parameters of our model, *D* and ρ, to specific patients, we consider the model in its spherically symmetric form. For data, we restrict ourselves to what can reasonably be expected to exist for each patient clinically, namely pretreatment T1Gd and T2 MRIs from two time points. We further make the assumption that the T1Gd and T2 regions of abnormality capture all regions with tumor cell density greater than or equal to 80% or 16% of the carrying capacity respectively. By converting the measured volumes of the abnormalities seen on the pretreatment images, we are able to directly estimate a radial velocity for individual patients. Further, we have developed a method leveraging the analytic expression for *D*/ρ derivable from the linearized version of (1) to estimate an individual’s *D*/ρ requires the T1Gd and T2 spherically equivalent radii measurements from the same day [13]. With these two equations involving two unknowns, one can solve for an individual’s *D* and ρ.

This is a specific example of how one might calibrate parameters in a PDE which utilizes specific information about behavior of the model. If such information is not available, there are still numerous ways to calibrate the parameters all stemming from the concept, “Find the parameters that minimize the error between the model predicted data points and the observed data points.” Thus, calibration problems often fall into the category of minimization problems. There are many algorithms to solve such problems such as steepest descent and newton-like methods [14], [15]. Of these two algorithms, steepest descent is based on first derivatives of the objective function and is so easier to implement but converges more slowly. In contrast, Newton-like algorithms utilize both the first and second derivatives of the objective function leading to faster convergence, though they are more difficult to implement. Both methods, however, are subject to finding local minimum rather than the global minimum when dealing with nonlinear problems.

